# The roles of DDR2 and substrate stiffness on cancer cell transcriptome and proliferation

**DOI:** 10.1101/2023.10.29.564363

**Authors:** Theadora Vessella, Steven Xiang, Cong Xiao, Madelyn Stilwell, Jason Shohet, Esteban Rozen, Susan Zhou, Qi Wen

**Affiliations:** Department of Chemical Engineering, Worcester Polytechnic Institute, 100 Institute Rd Worcester, MA 01609, USA; Bancroft School, 110 Shore Drive, Worcester, MA 01605, USA; Nash Family Department of Neuroscience, Friedman Brain Institute, Icahn School of Medicine at Mount Sinai, New York, NY 10029, USA; Black Family Stem Cell Institute, Icahn School of Medicine at Mount Sinai, New York, NY 10029, USA; Department of Biomedical Engineering, Wichita State University, 1845 Fairmount St, Wichita, KS 67260, USA; University of Massachusetts Medical School, Department of Pediatrics, 55 Lake Ave North, Worcester Massachusetts, 01566, USA; Crnic Institute Boulder Branch, BioFrontiers Institute, University of Colorado Boulder, 3415 Colorado Avenue, Boulder Colorado, 80303, USA; Department of Physics, Worcester Polytechnic Institute, 100 Institute Rd Worcester, MA 01609

**Keywords:** DDR2, extracellular matrix, collagen, RNA-seq, biomechanics, transcriptome, pro-proliferation, senescence, traction force

## Abstract

The interactions between cancer cells and the ECM regulate carcinogenesis. The collagen receptor kinase DDR2 is dysregulated in certain cancer cells, but its precise role in these malignancies remains unclear. In this study, we perform RNA-seq to determine how DDR2 and the biomechanical environment regulate cancer cell behaviors. We show that DDR2 knockdown in SH-SY5Y neuroblastoma cells inhibits proliferation and promotes senescence by regulating relevant genes. Increasing substrate stiffness reduces proliferation and promotes cell spreading but does not change senescence or transcriptome. Furthermore, DDR2 knockdown modulates cellular responses to substrate stiffness changes, unraveling a crosstalk between DDR2 and mechanosensing. These findings indicate DDR2 and ECM biomechanics regulate cancer cell behavior through distinct mechanisms, providing new mechanistic insights of cancer progression.

## Introduction

A hallmark of cancer cells that distinguishes them from normal differentiated cells is their unrestricted proliferative ability. Numerous genetic, signaling, and metabolic pathways have been implicated in cancer proliferation. Broadly, these factors are divided into pro-proliferative and anti-proliferative ones (1). The former includes those that promote DNA and protein synthesis including PI3 kinase and Ras-MAPK signaling, chromosome segregation, and telomere extension just to name a few, whereas the latter acts as a brake for proliferation including the tumor suppressors PTEN and p53 (2, 3). Genetic mutation of both positive and negative regulators has been causally linked to various cancer types. Not surprisingly, hampering cancer proliferation is a primary target of cancer drugs, for example, paclitaxel (Taxol), the most prescribed drugs for treating cancers, acts by inhibiting microtubule assembly to prevent cell division (4). Despite extensive studies, however, how cancer cells could escape from cell division regulation is not fully understood.

Cancer cells grow in a microenvironment wherein they closely interact with the extracellular matrix (ECM). As a major ECM component, collagen composition regulates various steps of cancer progression including growth, invasion, and metastasis, partly through activation of its canonical receptor integrin to regulate cytoskeleton organization and cell motility (5–7). Recently, discoidin domain receptor tyrosine kinase 2 (DDR2), a non-typical collagen receptor that is dysregulated in various cancer types, has emerged as a key signaling molecule in carcinogenesis (8, 9). Collagen binding to DDR2 activates its tyrosine kinase activity to initiate canonical pathways such as ERK/MAPK and PI3K/AKT signaling cascades (10–12). Despite these studies, how DDR2 regulates cancer cell behavior is incompletely understood. Besides providing biochemical cues that elicit signaling in cancer cells, ECM components also establish the biomechanical environment that critically controls cancer progression (13, 14). Upregulated collagen production and altered collagen fiber organization result in stiffening of the tumor ECM environment. Such increased tissue stiffness has been exploited as a marker for detection of solid tumors (15). Integrin-mediated mechanotransduction has been proposed to regulate ECM stiffness-dependent signaling pathways that play pivotal roles in cell growth, proliferation, and survival (16, 17). High ECM stiffness has been reported to facilitate cancer metastasis by triggering epithelial-mesenchymal transition (17, 18), promote cancer cell proliferation, and boost resistance to chemotherapy (19). A recent study further suggested that DDR2 regulates the integrin mediated mechanotransduction functions of cancer associated fibroblasts (20). However, whether DDR2 and ECM biomechanics could interact to regulate cancer cell behavior is yet to be determined.

To systematically examine the effects of DDR2 signaling and substrate stiffness on cancer cells, in the present study, we performed RNA-seq analysis of a human neuroblastoma cell line SH-SY5Y. This cell line has been extensively used as a model to study cancer progression (21). Moreover, because SH-SY5Y cells could be differentiated into a dopaminergic neuronal lineage, SH-SY5Y cells have also been used to investigate cell biology of dopaminergic neurons and Parkinson’s disease (22). We found that shRNA knockdown of DDR2 alters global gene expression of SH-SY5Y cells and inhibits cell proliferation. Moreover, we found that increasing substrate stiffness also slows down proliferation, similar to DDR2 knockdown. However, RNA-seq revealed no gene expression changes associated with increasing substrate stiffness. These data suggest that DDR2 signaling and biomechanics, two downstream effectors of ECM, could regulate cancer cell proliferation through different mechanisms, with or without involvement of gene expression.

## Results

### Bulk RNA-seq revealed a profound impact of DDR2 on transcriptome

To survey the effects of DDR2 knockdown on SH-SY5Y cells, we compared shCTRL and shDDR2 cells grown on collagen-coated 2 kPa Polyacrylamide (PAA) gels. SH-SY5Y cells were stably transduced with a lentiviral shRNA construct to target DDR2 expression (shDDR2 SH-SY5Y) or a non-targeting vector (shCTRL SH-SY5Y) (Figure 1A, see methods). We harvested the cells cultured on the PAA gels 24 hours after plating, extracted RNA, constructed libraries, did the sequencing, and analyzed the RNA-seq data (Figure 1B). Three biological repeats were conducted for each sample. Volcano plot and MA plot analysis of the total 3,923 differentially expressed genes (DEGs) revealed that 1,982 genes reduced their expression in shDDR2 cells, when compared to shCTRL, whereas another 1,941 genes upregulated their expression (Figure 2A-2B, Table S2). As expected, DDR2 was among the most significant reduced genes, with a 63% reduction in shDDR2 cells when compared to shCTRL cells with a p-adj value of 3.3 × 10^−35^(Figure 2A, S1A). Reduction of the DDR2 mRNA level was further confirmed by quantitative PCR (Figure S1B). Heat map and principal component analysis (PCA) revealed that shDDR2 and shCTRL groups were well segregated (Figure 2C-2D). With the number of DEGs representing ∼15% of the entire human genome, our results demonstrate a profound role of DDR2 in regulation of SH-SY5Y cell transcriptome.

**Figure 1.**
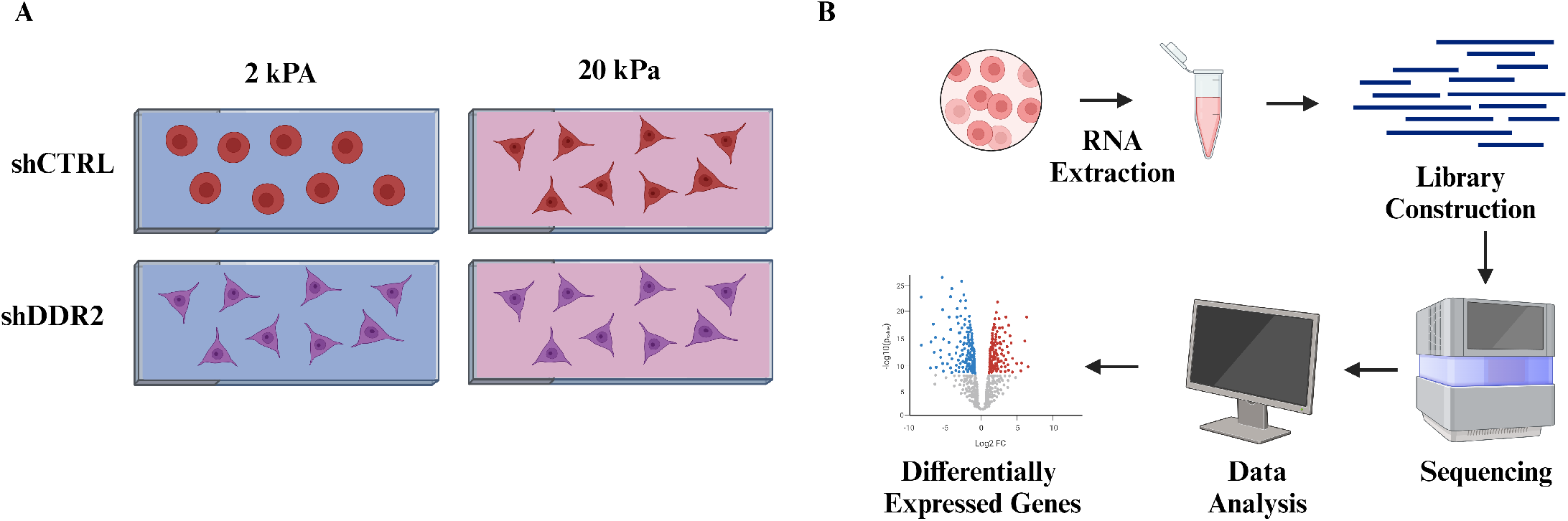
Schematic illustration of experimental process to find novel genes involved in biophysical stimulation. A) The four sample conditions used in this study: shCTRL and shDDR2 cells cultured on 2 kPa and 20 kPa collagen-coated polyacrylamide gel substrates. B) Experimental process of RNA-seq: a 24-hour incubation of the cells on PAA gels followed by cellular RNA extraction, library construction, sequencing, and data analysis. This figure is created with BioRender.com.

**Figure 2.**
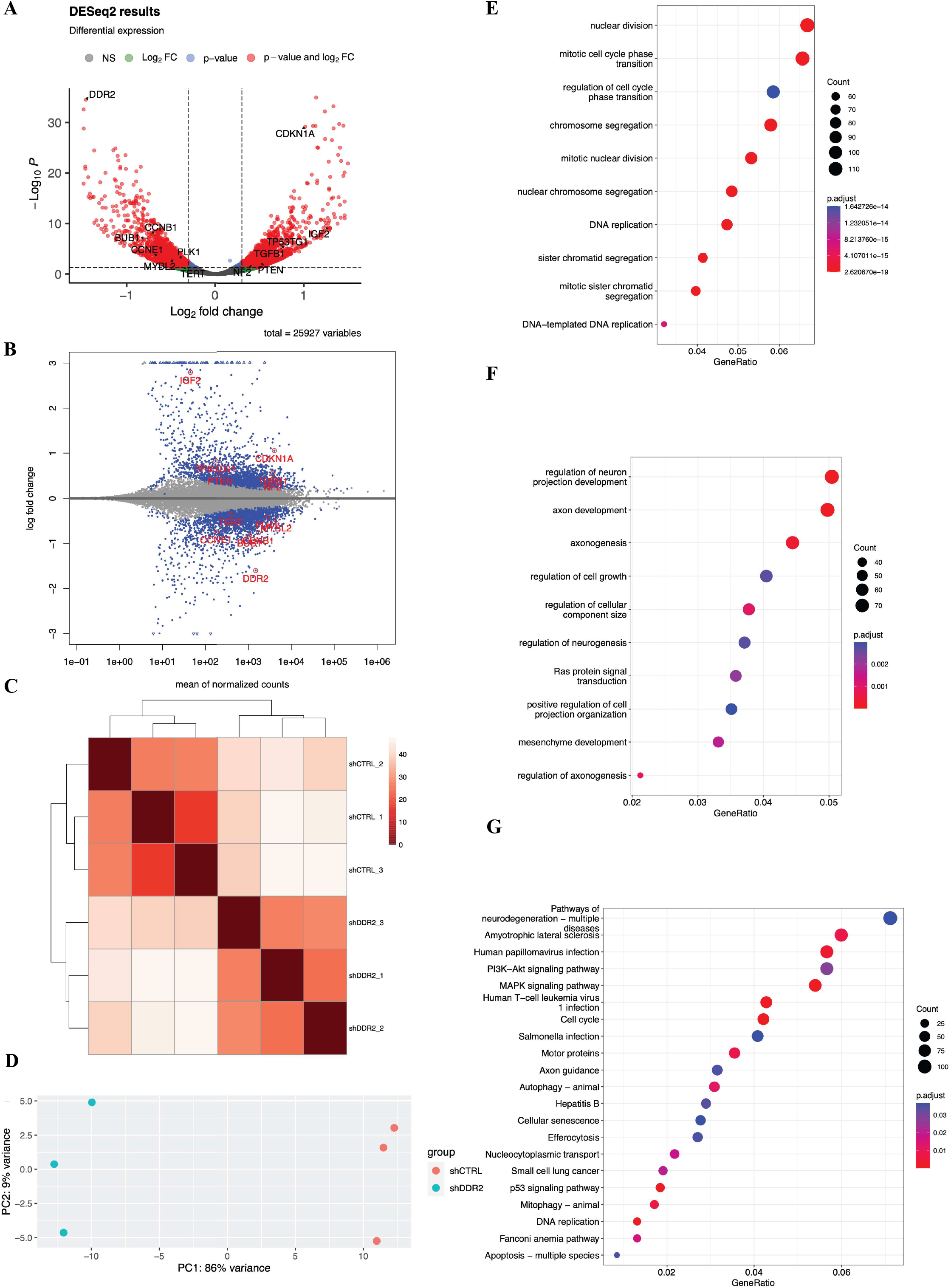
RNA-seq data analysis of shCTRL vs shDDR2 cells on soft substrates. (A) Volcano plot of RNA-seq data in the malignant neuroblastoma cell line SH-SY5Y where the x-axis represents fold change in transcripts from shCTRL vs shDDR2 cell lines (a positive score represents enrichment; a negative score represents depletion). The y-axis represents statistical confidence for each x-axis point. (B) MA plot of RNA-seq data, where the x-axis represents statistical confidence for each y-axis point. The y-axis represents fold change in transcripts from shCTRL vs shDDR2 cell lines. (C) Heatmap analysis of relationships among different samples. (D) PCA analysis of sample clustering. (E) GO biological process analysis of the down differentially expressed genes. (F) GO biological process analysis of the up differentially expressed genes. (G) KEGG enrichment analysis of the differentially expressed genes.

### Categorization and GO analysis

We performed gene ontology (GO) analysis of DEGs in shDDR2 SH-SY5Y cells. Strikingly, the GO biological process analysis showed that those down-regulated genes were enriched in the pathways related to cell proliferation, including nuclear division, mitotic cell cycle phase transition, chromosome segregation, DNA replication, and sister chromatid segregation, just to name a few (Figure 2E). Our analysis suggested that a normal function of DDR2 is to maintain fast proliferation of SH-SY5Y cells. When DDR2 is knocked down, proliferation of SH-SY5Y cells is expected to be reduced.

Next, we performed GO biological process analysis of those up-regulated DEGs in shDDR2 SH-SY5Y cells. Intriguingly, we found the top pathways were related to regulation of cell growth and regulation of cellular component size (Figure 2F), suggestive of stimulation of cellular growth and size after DDR2 knockdown. In addition, we noted that genes involved in neuronal development such as regulation of neurogenesis, axonogenesis, and regulation of neuron projection development were also upregulated (Figure 2F), suggesting differentiation of SH-SY5Y cells towards a neuronal fate after DDR2 knockdown. Indeed, SH-SY5Y cells are a neuroblastoma line and could be readily induced into dopaminergic neurons upon proper stimulation (21).

We further continued the pathway analysis by examining all DEGs, including up- and down-regulated genes using the Kyoto encyclopedia of genes and genomes (KEGG) database. One pathway showing significant gene enrichment was related to cell senescence, which is known as a viable but non-proliferation cell state (Figure 2G). Collectively, our GO and KEGG analyses support the notion that a normal function of DDR2 would be to maintain a fast proliferative, cancerous state of SH-SY5Y cells by preventing their differentiation.

### Identification of a gene cohort controlling proliferation, senescence, and cell size

The proliferative signature of cancer cells is known to be causally linked to expression of a core set of pro-proliferative genes including *MYBL2*, *BUB1*, and *polo-like kinase 1* (*PLK1*) (23). In addition, genes that promote cell cycle progression such as *CCNE1* and *CCNB1*, which encode cyclin E1 and cyclin B1, respectively, are also crucial players in cancer cell proliferation (23). Remarkably, we found a reduction of all these positive regulation genes of cell proliferation in shDDR2 SH-SY5Y cells (Figure 2A-2B, S2A).

As mentioned above, cell proliferation is also subject to negative regulation. For example, *phosphatase and tensin homolog* (*PTEN*) is one of best-known tumor suppressors. In human tumors, *PTEN* expression is usually suppressed through promoter methylation, allowing cancer cells to proliferate unrestrictedly. As expected, we found that *PTEN* expression was significantly increased after DDR2 knockdown (Figure S2B). Consistently, expression of *NF2*, another tumor suppressor (24), was increased after DDR2 RNAi (Figure S2B). These data suggested that DDR2 knockdown could reduce the proliferative competence. Along this line. *TGF-β* is known for its antiproliferative effects (25), and *TGF-β* expression was increased after DDR2 knockdown (Figure 2A-2B, S2B).

After each cell division, the length of telomere is reduced and such reduction is thought to limit or prevent normal cells from unlimited proliferation (26). In normal cells, expression of an enzyme known to increase the telomere length, *telomerase reverse transcriptase* (*TERT*), is largely suppressed. However, cancer cells are unique in that they could upregulate *TERT* expression to ensure telomere length increases after each cell division (26). We noted that *TERT* expression was significantly reduced after DDR2 knockdown (Figure S2C). Such *TERT* reduction has been linked to cancer senescence (27). Supporting the notion that shDDR2 SH-SY5Y cells are poised to enter senescence, a signature senescence gene, *CDKN1A* (28), was upregulated after DDR2 knockdown (Figure S2C). Moreover, increasing cell growth and size is often considered as a morphological marker of cell senescence. Indeed, we found that *insulin growth factor 2* (*IGF2*), which is known to stimulate cell growth (29), was increased after DDR2 knockdown (Figure 2A-2B, S2C).

### Bulk RNA-seq revealed no transcriptome changes in response to substrate stiffness

Cancer progression is often associated with changes of their biomechanical properties (14). Indeed, hardness is often the first sign of tumorigenesis. Moreover, the biomechanical environment surrounding cancer cells actively regulates almost all steps of cancer progression (13). Despite the close interactions between cancer cells and the surrounding biomechanical cues, the impact of biomechanics on cancer cell transcriptomes is yet unknown. Therefore, we performed bulk RNA-seq to profile SH-SY5Y cells grown on 2 kPa and 20 kPa substrate stiffness. Surprisingly, we found no statistically significant changes in any gene expression across the transcriptome (Figure 3A-3B, 3E). Heatmap and PCA analysis showed that these two groups cannot be separated (Figure 3C-3D). We also explored DEGs in shDDR2 SH-SY5Y cells grown on 2 kPa and 20 kPa substrate stiffness and found no statistically significant changes of gene expression (data not shown). Nevertheless, we did notice that DEGs associated with DDR2 knockdown were different when cells were cultured at 2 kPa (3,923 DEGs) versus 20 kPa (1,013 DEGs) (Table S2, S3).

**Figure 3.**
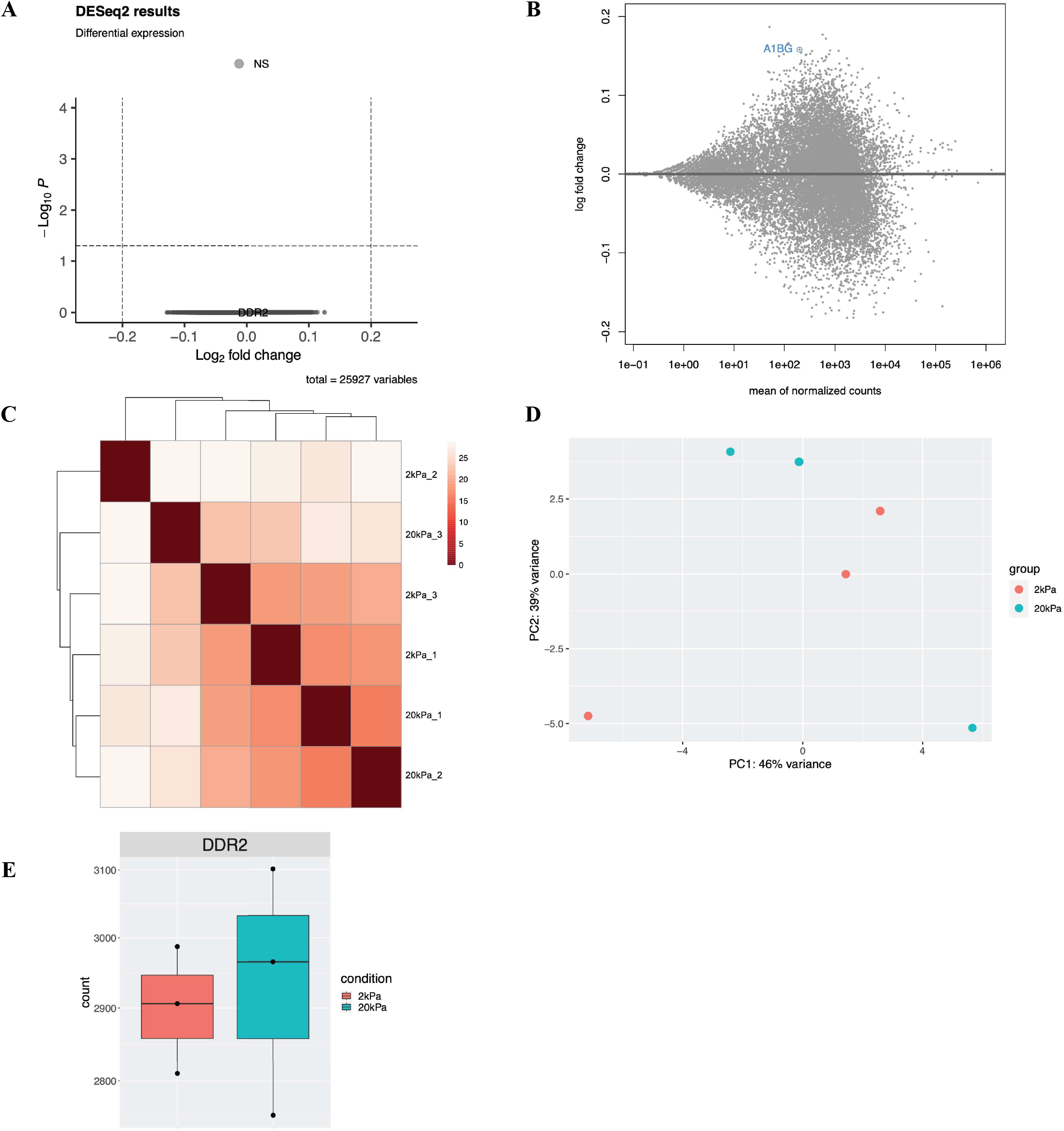
RNA-seq data analysis of shCTRL cells cultured on hard vs soft substrates. (A) Volcano plot of RNA-seq data in the malignant neuroblastoma cell line SH-SY5Y, where the x-axis represents fold change in transcripts from shCTRL cells cultured on hard vs soft substrates (a positive score represents enrichment, a negative score represents depletion). The y-axis represents statistical confidence for each x-axis point. (B) MA plot of RNA-seq data, where the x-axis represents statistical confidence for each y-axis point. The y-axis represents fold change in transcripts from shCTRL cell lines cultured on hard vs soft substrates. (C) Heatmap analysis of relationships among different samples. (D) PCA analysis of sample clustering. (E) The normalized reads count of shDDR2 in the shCTRL cell line cultured on hard vs soft substrates.

### The effects of DDR2 knockdown and substrate stiffness on cell proliferation and senescence

Our RNA-seq data predicted a reduction of cell proliferation after DDR2 knockdown. To directly test this idea, we performed the EdU cell proliferation assay. EdU as an analog of thymidine is incorporated into DNA selectively in dividing cells. We found that when grown on substrates of either 2 kPa or 20 kPa, EdU labeling was significantly reduced in shDDR2 when compared to shCTRL SH-SY5Y cells (Figure 4A). These results provide direct evidence of reduced proliferation after DDR2 knockdown, thus validating our RNA-seq results.

**Figure 4.**
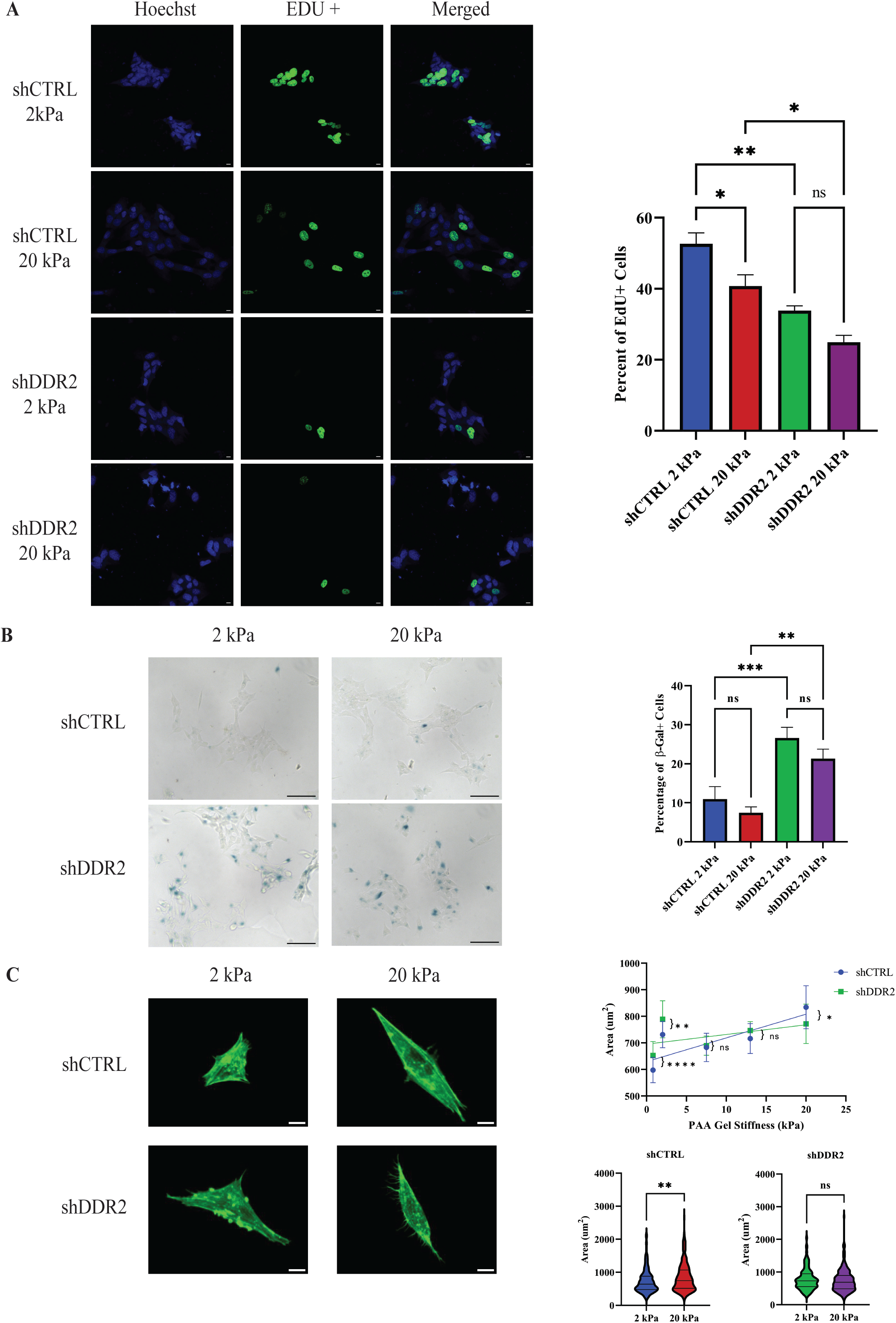
DDR2 knockdown leads to senescence and increased cell area in shDDR2 neuroblastoma cells. A) Representative images of EdU assay on shCTRL and shDDR2 cells for both soft and hard gels. Blue channel is Hoechst 33342 and green is the positive EdU signal. Percentage of positive EdU cells taken as ratio of EdU positive cells / total number of cells (n = 6-15 regions). One-way ANOVA followed by Bonferroni post-hoc test, not significant (ns) p>0.05, *p≤ 0.05, **p≤ 0.01, *** p≤ 0.001. B) Senescence analysis of shCTRL and shDDR2 cell lines using beta-galactosidase stain after 48 hours in culture. Blue arrow represents a positive beta-galactosidase signal. One-way ANOVA followed by Bonferroni post-hoc test, not significant (ns) p>0.05, *p≤ 0.05, **p≤ 0.01, *** p≤ 0.001. Percentage of positive beta-galactosidase cells taken as ratio of beta-galactosidase positive cells / total number of cells (n=10 regions). C) Representation of cell areas for shCTRL and shDDR2 cells on soft and hard PAA gel (N = 3-4, n = 224-261 cells). Line graph represents the mean area as a function of substrate stiffnesses (0.8, 2, 7.5, 13, and 20 kPa) based on Mann-Whitney Test, not significant (ns) p>0.05, *p≤ 0.05, **p≤ 0.01, *** p≤ 0.001, ****p≤0.0001. All error bars represent s.e.m. and all scale bars represent 10 μm.

We also observed a significant reduction in EdU incorporation in shCTRL SHY-SY5Y cells when substrate stiffness was increased from 2 kPa to 20 kPa (Figure 4A). This suggests that an increase in substrate stiffness could reduce SH-SY5Y cell proliferation, despite our RNA-seq analysis showing no associated changes in gene expression (Figure 3A-3B). On the other hand, increasing of substrate stiffness failed to alter the proliferation of the shDDR2 SH-SY5Y cells (Figure 4A), which had down regulated DDR2 expression. These results suggest that DDR2 plays an important role in regulating cellular response to substrate stiffness.

Cell senescence is a common outcome of cancer cells that exit cell cycles (30). To measure cell senescence, we stained for the enzymatic activity of β-galactosidase, a marker of cell senescence (31). We found that β-galactosidase signals were significantly increased in shDDR2 SH-SY5Y cells when compared to shCTRL SH-SY5Y cells, when cultured at either 2 kPa or 20 kPa stiffness (Figure. 4B, S3). This result suggests that DDR2 knockdown induces senescence of SH-SY5Y cells, which is consistent with our RNA-seq analysis (Figure 2G). On the other hand, the change in substrate stiffness did not result in significant changes in the β-galactosidase signals in either shCTRL or shDDR2 SH-SY5Y cells (Figure 4B).

We further examined cell area as senescence is associated with increased cell size (32–34). The spreading areas of shCTRL cells and shDDR2 cells cultured on substrates of varying stiffness (from 0.8 to 20 kPa) were measured and plotted as a function of substrate stiffness. shCTRL cells increased their spreading area when substrate stiffness was increased (Figure 4C, S4). However, shDDR2 cells did not have an obvious increase in spreading area with increasing substrate stiffness (Figure 4C, S4), suggesting that DDR2 is indispensable for SH-SY5Y cells to respond to stiffness changes. Moreover, at lower stiffnesses (0.8-2 kPa), shDDR2 cells exhibited larger areas than shCTRL cells (Figure 4C), consistent with cells entering a senescence state. Taken together, our results indicate that downregulation of DDR2 in neuroblastoma cells decreases cell proliferation and induces senescence that is independent of substrate stiffness.

### Effects of substrate stiffness and DDR2 knockdown on cell contractility

Tumor cells generate force to remodel the ECM and facilitate metastasis. Cellular traction force has previously been shown to correlate with increasing metastatic potential of cancer cells (35–37). To investigate whether cancer cells in the fast proliferative versus senescence states could exhibit different traction force, we measured traction forces that shCTRL and shDDR2 cells exerted on substrates of varying stiffness (from 0.8 to7.5 kPa). We found that, either on soft substrates (0.8 kPa) or on hard substrates (7.5 kPa), shCTRL cells exhibited a greater total force than shDDR2 cells (Figure 5A-5D). These results underline the importance of DDR2 in cell contractility and indicate that SH-SY5Y neuroblastoma cells in the fast-proliferating state are more contractile than those in the senescent state.

**Figure 5.**
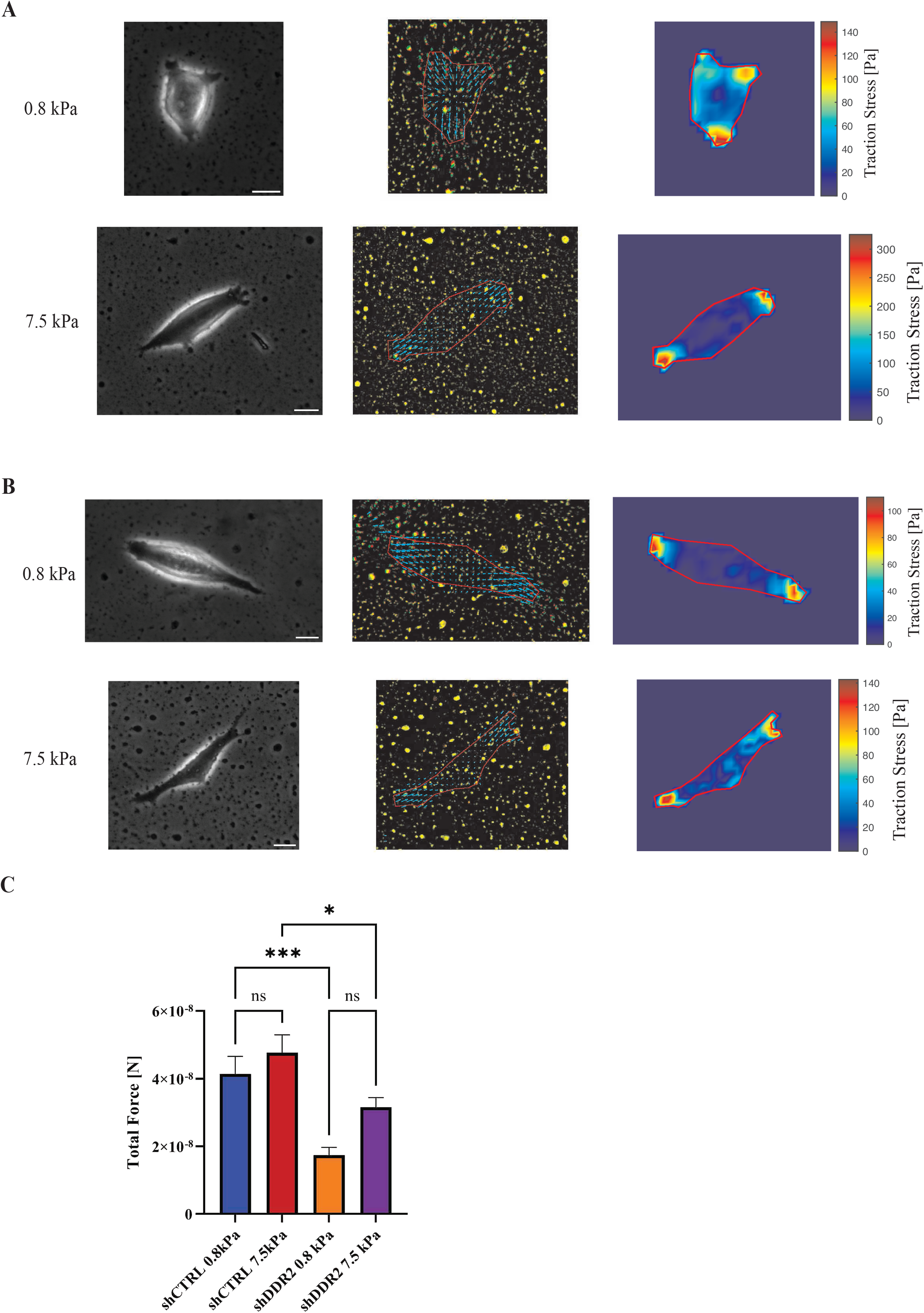
**Traction force microscopy on varying stiffness of collagen coated PAA gels for shCTRL and shDDR2 cell lines**. Representative phase contrast images, bead displacement maps, and stress heat maps for A) shCTRL and B) shDDR2 cell lines on 0.8 and 7.5 kPa collagen coated PAA gels. Results summary of (C) traction forces of shCTRL and shDDR2 on 0.8 kPa and 7.5 kPa substrate stiffnesses (n = 26-29 cells). Error bars represent s.e.m.. Scale bars represent 10 μm. One-way ANOVA with Bonferroni post-hoc test, not significant (ns) p>0.05*p≤ 0.05, **p≤ 0.01, *** p≤ 0.001.

## Discussion

The biomechanical properties of ECM change dynamically throughout cancer progression. The stiffened tumor ECM and abnormal mechanosensitivity of cancer cells have been shown to promote metastasis (38–40). The ECM receptors on cell surface are key components for transduction of these biochemical cues into intracellular signals. As a non-typical collagen receptor, DDR2 binds to fibrous collagen I. Dysregulated DDR2 expression has been documented in various cancer types including neuroblastoma. However, how neuroblastoma cells respond to substrate stiffness, as well as the role of DDR2 in sensing substrate stiffness are still poorly understood.

In the present study, we provide to our knowledge the first RNA-seq study to comprehensively measure the effects of substrate stiffness on neuroblastoma cells. We found no changes of gene expression between SH-SY5Y cells grown on a soft 2 kPa versus a hard 20 kPa substrate. This result is rather unexpected given that we have observed overt changes of cell proliferation, cell size, and cell contractility in response to stiffness changes. On the other hand, knocking down the collagen receptor DDR2 profoundly altered SH-SY5Y cell transcriptome. Pathway analysis revealed a down-regulation of pro-proliferative genes and an upregulation of tumor suppressor genes. These predictions were later validated by EdU labeling, β-galactosidase staining, and cell morphology assays.

Although the roles of DDR2 in the regulation of cell homeostasis, migration or cell cycle remain controversial, a growing number of reports have implicated DDR2 as a key driver of cell proliferation in fibroblasts (41, 42), chondrocytes (41, 43, 44), hepatic stellate cells (45), lung squamous cancer cell lines (46), osteoblasts (44), and breast cancer cell lines (47). Therefore, it is not surprising to observe that DDR2 is also required for proper neuroblastoma cell proliferation. Furthermore, recent work from Xu et al. (48) suggested that epigenetic down-regulation of DDR2 in human-derived bone marrow mesenchymal stem cells is associated with reduced proliferation and increased senescence of these cells, which is in agreement with our findings in SH-SY5Y neuroblastoma cells. Importantly, a previous work demonstrates that pharmacologic inhibitions or genetic ablation of DDR2 in several neuroblastoma cell lines impairs their proliferation and *in vivo* tumor growth and metastasis (Rozen et al., 2023 under revision).

The overlapped cellular responses between DDR2 knockdown and increasing of substrate stiffness, such as proliferation arrest, nevertheless suggest that genetic factors and biomechanical cues could elicit similar cellular responses in SH-SY5Y cells through distinct mechanisms, with or without involvement of gene expression. What are the possible cellular responses activated by substrate stiffness? Mechanosensing is known to trigger complex intracellular signaling (49). Indeed, ECM stiffness could affect multiple signaling pathways such as PI3K-AKT, YAP/TAZ, and Rho/ROCK (50). The typical response times of these pathways are normally in the range of minutes to hours (51). Earlier studies also show that it requires weeks of culture on a substrate for cells to alter their gene transcription (52, 53). Therefore, mechanosensing could act as the early phase response to ECM stiffness, while gene expression, which normally takes longer time to happen, could be a result of such signaling events through a feedback regulation.

shDDR2 SH-SY5Y cells partially lost their responses to substrate stiffness as demonstrated by the EdU and cell area assays. These findings support a role of DDR2 in modulating mechanosensing of SH-SY5Y cells. Providing that some signaling pathways such as MAPK and PI3K-AKT are regulated by both DDR2 and substrate stiffness (54), it is plausible that DDR2 modulates mechanosensing of substrate stiffness by regulating these pathways.

## Materials and Methods

### Data Availability

RNA-seq data have been deposited at GEO with the accession number GSE246550 and will be publicly available upon publication. All other data are available upon request to the corresponding author Dr. Qi Wen.

### Cell Culture

Human neuroblastoma cell line SH-SY5Y stably transduced with the Tet-pLKO-puro lentiviral vector expressing either a control non-targeting shRNA or shDDR2 were kindly provided by Dr. Jason Shohet (UMass Chan Medical School). Cells were cultured in Dulbecco’s modified Eagle’s medium (DMEM) supplemented with 10% fetal bovine serum (Gibco), 2mM glutamine (Gibco) and antibiotics (penicillin and streptomycin) (Gibco).

### Polyacrylamide Substrate Preparation

Polyacrylamide gel substrates were prepared through the polymerization of acrylamide and bis-acrylamide, with varying concentrations to achieve the desired stiffness levels (Table S1). This polymerization process was initiated by a solution containing 0.1% ammonium persulfate and 0.3% N,N,N’,N’-tetramethylethylenediamine. Collagen type I (Corning) was crosslinked to the PAA gel surface using sulfo-SANPAH (Proteochem). The gels were submerged under 1mg/ml sulfo-SANPAH (G-Biosciences, St. Louis, MO) solution and placed 2 inches below an 8 W ultraviolet UV lamp (Hitachi F8T5 – 365nm) and irradiated for 15 minutes. The gels were then washed with HEPES buffer and soaked with 0.1 mg/ml rat-tail collagen type I (Corning) for 12 hours at 4 °C. After collagen coating, the collagen was aspirated, and gels were placed in culture medium and incubated for 30 minutes at 37°C before cells were seeded on them.

For traction force microscopy, we followed an established protocol (55) to fabricate gel disks of 18 mm in diameter and approximately 100 μm in thickness. These gel disks were prepared with 0.1 μm red fluorescent beads (Life Technologies) embedded just beneath the top surface. 25 mm x 25 mm square glass coverslips (VWR) were cleaned, treated with 1% 3-aminopropyl-trimethoxysilane solution for 10 minutes, and then coated with 0.5% glutaraldehyde. Round glass coverslips (VWR), 18 mm in diameter, were plasma cleaned and coated with a thin layer of fluorescent beads. A 25 μL mixture of acrylamide, bis-acrylamide, and initiators was applied between the glutaraldehyde-coated square coverslip and the beads-coated round coverslip, followed by polymerization at room temperature for 15 minutes. Subsequently, the round coverslip was gently removed, leaving the resulting gel disk firmly attached to the square coverslip, with the embedded beads positioned within 2 μm below the gel surface. To allow for the increased cell numbers needed for RNA testing, PAA gels were prepared onto 75×50 mm microscope slides (Corning) according to previously described protocol (56).

### Lentivirus preparation and infection

HEK-293T cells were maintained at 37°C in Dulbecco’s modified Eagle medium (DMEM), supplemented with 10% FCS and antibiotics (100 units/ml penicillin and 100 μg/ml streptomycin). Cells were transfected with pVSV-G (57) and pCMVΔR8.91 (58), together with the pLKO.1-puro non-targeting vector (Sigma Mission clone SHC016; ‘shCTRL’) or the pLKO.1-shRNA vector (Sigma Mission TRCN0000001418; ‘shDDR2’) using Lipofectamine^TM^ 2000 reagent (Thermo Fisher Scientific) as recommended by the manufacturer and following the recommendations of the RNAi Consortium (TRC) laboratory protocols with slight modifications. Twelve hours after transfection the medium was replaced by DMEM, supplemented with 30% FCS and antibiotics which. Cell supernatants were harvested every 24 hours, replaced with fresh medium, and stored at 4°C until collection of the last harvest (at 72 hour). At this point, the consecutive harvests were pooled, filtered through 0.45mm filters and split in 3-5 mL aliquots, which were stored at −80°C. SH-SY5Y cells were infected with shCTRL or shDDR2 lentiviral particles by adding a 1:1 mix of medium:viral supernatant for 24-48 hours. Puromycin selection (2 μg/ml) was applied for 2-3 days and always compared to non-transduced control cells, which generally died within the first 24 hs. DDR2 downregulation was confirmed by Western Blot and PCR.

### RNA sequencing and data analysis

For the cell samples from different genotypes or conditions, RNA was extracted using the RNeasy microKit (QIAGEN) combined with DNase (TurboDNase) (ThermoFisher). The following RNA concentration and integrity were measured by the Qubit 4 Fluorometer (Thermo Fisher Scientific) and 2100 Bioanalyzer (Agilent), respectively. We used the NEBNext® Ultra™ II Directional RNA Library Prep Kit for Illumina® kit to make all RNA libraries without any modifications. For each library, the input of approximately 500 ng total RNA, and a total of nine PCR cycles were used for library amplification. The quality and concentration of libraries were measured by 2100 Bioanalyzer (Agilent) and Qubit 4 Fluorometer (Thermo Fisher Scientific), respectively. All libraries were normalized to 20 nM and pooled together, the final concentration of library pool was 5 nM, loaded onto the NextSeq500/550 High Output Kit v2.5 (150 Cycles) chip (Illumina), and sequenced on the NextSeq550 (Illumina). Reads were sorted by the barcodes assigned to each library and adapter sequences were removed using Trimmomatic (59). The reads were then mapped to the human genome through STAR (60), and the gene expressions count table was generated by Tetranscripts (61). DEGs were identified using a R package DeSeq2 (62). GO analysis was performed using another R package clusterProfiler (63).

### RT-quantitative PCR

RNA was extracted as described above and 100 ng of RNA was reverse transcribed into cDNA using a Superscript IV kit (ThermoFisher). The RT-quantitative PCRs were carried out in four biological replicates in a 96-well plate (QIAGEN) using a QIAcuity system (QIAGEN). A QIAcuity EvaGreen (EG) PCR Kit (QIAGEN) was used with gene-specific primers for DDR2 and GAPDH. All transcript levels were normalized to the GAPDH transcript level.

### Cell Senescence Analysis

Cells were plated onto collagen coated PAA gels of desired stiffness and allowed to incubate for 48 hours. Medium was removed from cells after 24 hours. Cells were rinsed with pre-warmed 1x PBS and fixed with 1x fixative solution provided by senescence beta-galactosidase staining kit (9860, Cell Signaling Technology) for 15 minutes. Fresh beta-galactosidase staining solution was prepared according to manufacturer’s instructions and pH was confirmed to be 6.0. Cells are washed 2x with 1x PBS and 3mL of staining solution are added to each dish. Dishes are wrapped in parafilm to avoid evaporation and incubated at 37°C in a dry incubator until blue color was visible (48 hours). The beta-galactosidase positive cells (blue) were considered as positive senescent cells.

### Cell Proliferation Assay

Cell proliferation was assessed using, ethynyl-2′-deoxyuridine (EdU) incorporation using Click-iT EdU imaging kit (Thermo Fisher). Cells were seeded onto collagen coated PAA gels of varying stiffnesses for 24 hours. Following overnight incubation and relaxation of cells, half of the cell media was removed and replaced with 20 μM EdU working solution and allowed to incubate in ideal culture conditions for 4 hours. After the 4 hours of EdU labeling, cells are fixed using 3.7% formaldehyde in PBS for 15 minutes at room temperature. After fixation, cells are rinsed 2x with 3% BSA in PBS. The washing solution was removed, and cell membranes were permeabilized with 0.5% Triton-X in PBS incubated at room temperature for 20 minutes.

Permeabilization solution was removed, and dishes are rinsed 2x with 3% BSA in PBS. Reaction cocktail was prepared fresh and according to the manufacturer’s protocol and incubated on each dish at room temperature for 30 minutes, protected from light. Reaction cocktail is removed, and dishes are washed once with 3% BSA in PBS. Hoechst 33342 is diluted at 1:2000 in PBS and incubated in each dish for 30 minutes at room temperature, protected from light. Dishes were rinsed 2x with PBS. Final dishes were imaged in PBS. EdU and Hoechst 33342 signals were captured in separate channels using a Zeiss LSM 700 confocal microscope with a water immersion 20x 1.0 NA objective, and maximum projection images were processed with ImageJ software (NIH). Microscopy settings were held constant between experiments. Green EdU signal was considered as a positive EdU signal.

### Phalloidin Imaging

Cells were incubated onto gels for 24 hours; pre-warmed PBS were used to rinse cells 3 times. After washing, cells were immediately fixed using a 3.7% formaldehyde for 15 minutes at room temperature. Cells are washed twice with PBS, and cell membranes are permeabilized using a 0.1% Triton-X solution for 20 minutes. After Triton-X incubation, cells are washed 2 times with PBS. To stain F-Actin within cells, 0.5 μL of 400x stock of Phalloidin Alexa 488nm (Thermo Fisher) is added per 200 μL of PBS to each sample and incubated at room temperature away from light for 1 hour. Cells are then washed 2x PBS. Final dishes were imaged in PBS under Zeiss LSM 700 confocal microscope at 20x 1.0 NA and Z-stacks were taken from the bottom to top of individual cells. Microscopy settings were held constant between experiments. Average projections of z-stacks were created within ImageJ.

### Cellular spreading area measurements

Cells were seeded with 20% confluency on PAA gels for 24 hours. Individual cells were imaged using an Olympus IX83 inverted microscope equipped with a 40x 0.6 NA objective using the Phase contrast mode. Cell area was measured from the phase contrast images using Fiji ImageJ software (NIH). Cell boundaries were traced using the free-hand tool and the area enclosed by the cell boundary was measured as cell area.

### Cell traction force measurements

Cell traction forces were measured using traction force microscopy (64). Cells were cultured on PAA substrates for 24 hours before subjected to traction force microscopy. For each cell selected for traction force microscopy, a fluorescence image of the substrate was recorded to capture the marker beads in the stressed state. In addition, a phase-contrast image was acquired to record the morphology of the cell. Trypsin (Invitrogen) was then applied to disrupt cell-substrate interactions and cause the cell to detach from the substrate. A final fluorescent image of the substrate was taken to capture the marker beads in the relaxed, unstressed state. Bead displacements were calculated from the two fluorescent images using a particle image velocimetry toolbox written in MATLAB (65). Traction stress on the gel surface was calculated from the bead displacements using the finite element analysis software (Ansys, Inc). The magnitude of total traction force (F) was calculated by integrating the magnitude of traction stress over the cell area. Bead displacements were calculated from the two fluorescent images using a particle image velocimetry toolbox written in MATLAB (65). Traction stress on the gel surface was calculated from the bead displacements using the finite element analysis software (Ansys, Inc). The magnitude of total traction force (F) was calculated by integrating the magnitude of traction stress over the cell area.

## Author Contributions

TV, SX, MS, QW and SZ conceived and designed the experiments. TV, SX and MS prepared the experimental materials. TV, SX, MS, and ER performed the experiments. TV, SX, MS, CX, ER, QW and SZ interpreted and analyzed the data. TV, SX, CX, ER, SZ and QW wrote the manuscript.

## Acknowledgements

The work of Madelyn Stilwell is supported by the NSF REU Site grant 2150076.

