## Supplemental Information for "The roles of DDR2 and substrate stiffness on cancer cell transcriptome and proliferation"

### SUPPLEMENTAL FIGURES

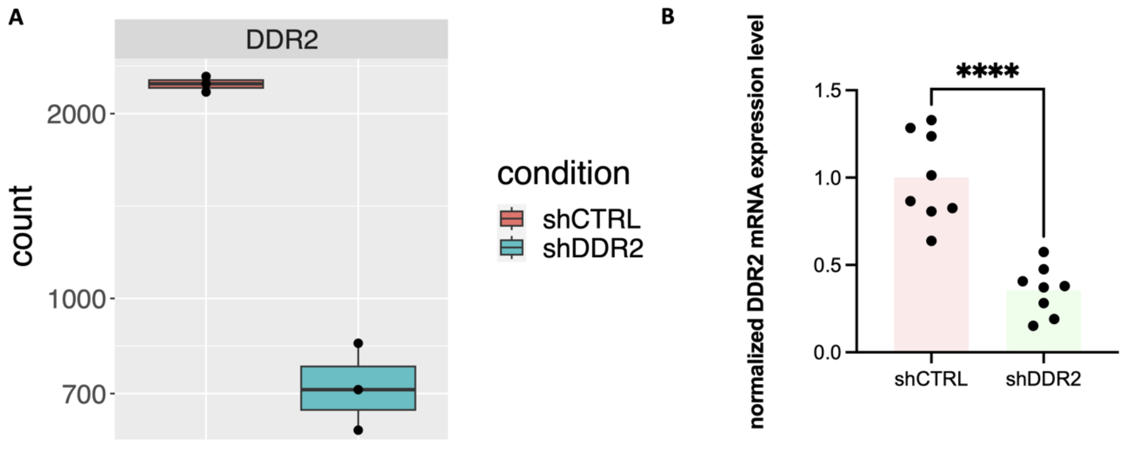


**Figure S1.** DDR2 downregulation from A) RNA seq and validation by B) q-PCR between shCTRL and shDDR2 cell lines.

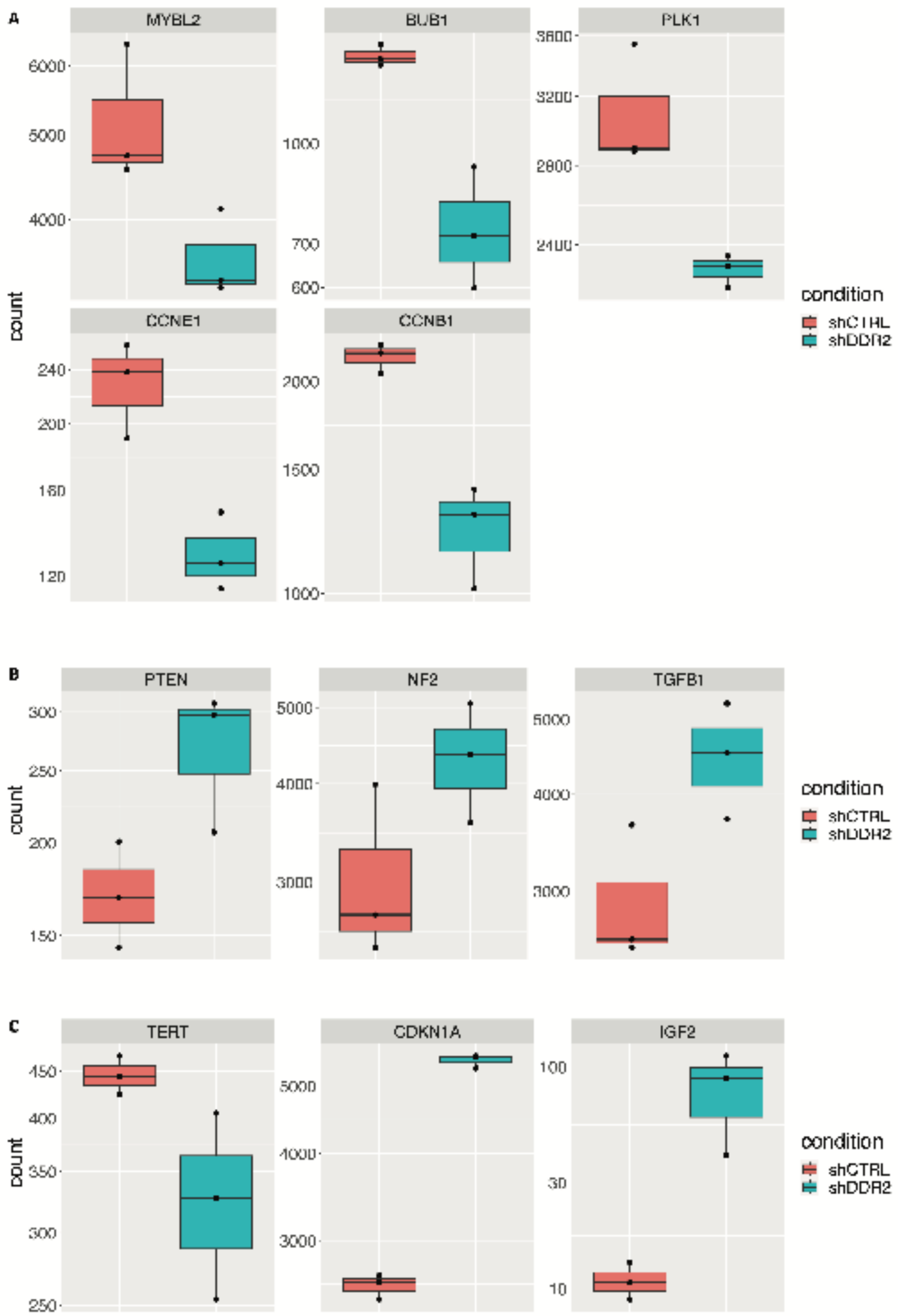


**Figure S2. The normalized sequencing reads counts of genes involved in cell cycle and cellular senescence pathways from shCTRL vs shiDDR2 cell lines**. (A) The normalized reads count of *MYBL2*, *BUB1*, *PLK1*, *CCNE1* and *CCNB1* in shCTRL vs shiDDR2 cell lines. (B) The normalized reads count of *PTEN*, *NF2* and *TGFβ1* in shCTRL vs shiDDR2 cell lines. (C) The normalized reads count of *TERT, CDKN1A* and *IGF2* in shCTRL vs shiDDR2 cell lines.

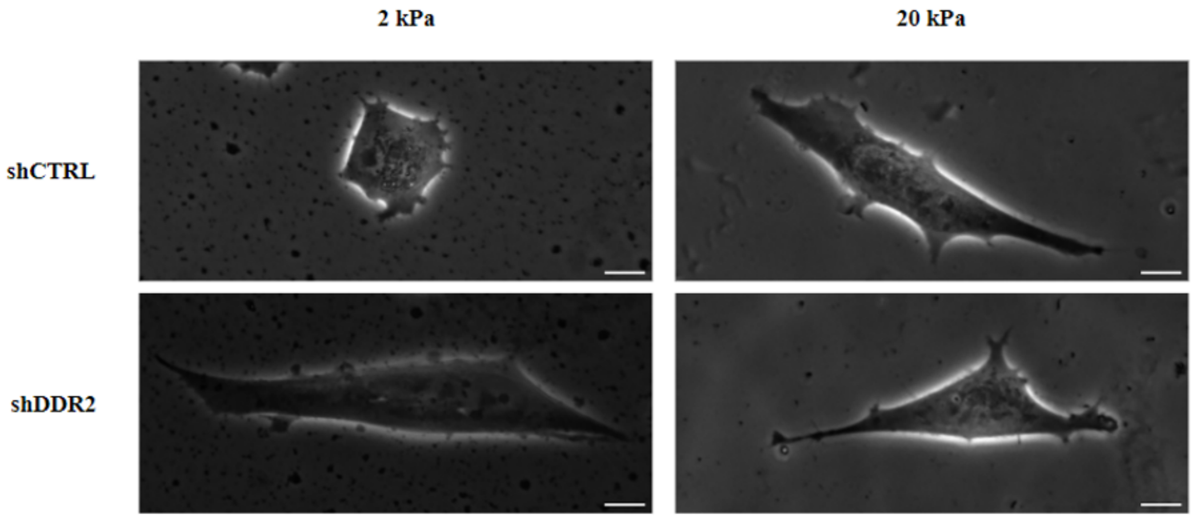


**Figure S3.** Representative phase contrast images of cellular morphology of shCTRL and shDDR2 on both 2 kPa and 20 kPa PAA gels. Scale bars represent 10 $\mu$m.

**Table S1. Stiffnesses of PAA Gels**

| Stiffness (kPa) | % ACL | % BIS |
|---|---|---|
| 0.8 | 2 | 0.12 |
| 2 | 5 | 0.08 |
| 7.5 | 8 | 0.08 |
| 13 | 10 | 0.12 |
| 20 | 12 | 0.14 |
